# Drp1-mediated mitochondrial fission regulates calcium and F-actin dynamics during wound healing

**DOI:** 10.1101/727628

**Authors:** Susana Ponte, Lara Carvalho, Maria Gagliardi, Isabel Campos, Paulo J. Oliveira, António Jacinto

## Abstract

Mitochondria adapt to cellular needs by changes in morphology through fusion and fission events, referred to as mitochondrial dynamics. Mitochondrial function and morphology are intimately connected and the dysregulation of mitochondrial dynamics is linked to several human diseases. In this work, we investigated the role of mitochondrial dynamics in wound healing in the *Drosophila* embryonic epidermis. Mutants for mitochondrial fusion and fission proteins fail to close their wounds, indicating that the regulation of mitochondrial dynamics is required for wound healing. By live-imaging, we found that loss of function of the mitochondrial fission protein Dynamin-related protein 1 (Drp1) compromises the increase of cytosolic and mitochondrial calcium upon wounding and leads to F-actin defects at the wound edge, culminating in wound healing impairment. Our results highlight a new role for mitochondrial dynamics in the regulation of calcium and F-actin during epithelial repair.

**Summary:** We show that mitochondrial dynamics proteins are required for epithelial repair. In particular, Drp1 loss-of-function leads to defects in the dynamics of cytosolic and mitochondrial calcium and F-actin upon wounding.

## Introduction

Mitochondria perform critical cellular functions such as energy production, regulation of calcium (Ca^2+^), redox homeostasis and cell death (El-Hattab and Scaglia, 2016). Mitochondrial shape is controlled by antagonizing fusion and fission events (Lewis and Lewis, 1914; Nunnari et al., 1997), described as mitochondrial dynamics, that allow mitochondria to adapt to cellular demands (Nunnari and Suomalainen, 2012).

Dynamin-related proteins regulate mitochondrial dynamics through their GTPase activity (Hoppins et al., 2007). Mitochondrial fission is accomplished by Dynamin-related protein 1 (Drp1). Upon activation, Drp1 is recruited from the cytosol to mitochondria, oligomerizes and constricts this organelle until division is achieved (Bleazard et al., 1999; Labrousse et al., 1999; Smirnova et al., 2001; Yoon et al., 2001). Mitochondrial fusion requires the merge of both the outer (OMM) and the inner mitochondrial membranes (IMM). Mitofusin 1 (Mfn1) and Mitofusin 2 (Mfn2) are responsible for OMM fusion (Rojo et al., 2002), while Optic atrophy 1 (Opa1) mediates fusion of the IMM (Griparic et al., 2004; Olichon et al., 2003).

Regulation of mitochondrial dynamics is essential for development (Chen et al., 2003; Ishihara et al., 2009; Waterham et al., 2007) and dysregulation of its machinery is implicated in a wide range of human diseases, including neuropathies, type II diabetes, and cancer (Anderson et al., 2018; Ranieri et al., 2013; Rovira-Llopis et al., 2017). However, the role of mitochondrial dynamics in other contexts, such as epithelial repair is still largely unknown.

Wound healing in simple epithelia is characterized by the accumulation of F-actin and Non-muscle Myosin II (Myosin) at the cell boundaries that face the wound, forming an actomyosin cable that contracts and brings cells together, thereby closing the wound (Bement et al., 1999; Danjo and Gipson, 1998; Kiehart et al., 2000; Xu and Chisholm, 2011). Additionally, wound healing involves cell crawling mediated by actin protrusions (Abreu-Blanco et al., 2012; Verboon and Parkhurst, 2015) and cellular rearrangements (Carvalho et al., 2018; Razzell et al., 2014).

Recent studies have addressed the role of mitochondria in tissue repair, showing that mitochondrial reactive oxygen species (ROS) promote wound healing by regulating F-actin and Myosin at the wound edge, either by acting on Rho GTPases (Muliyil and Narasimha, 2014; Xu and Chisholm, 2014) or on cell-cell junction remodeling (Hunter et al., 2018). In this work, we show that the mitochondrial dynamics machinery is required for repair, as mutants for these proteins fail to close epithelial wounds. In particular, the fission protein Drp1 is required for F-actin accumulation at the wound edge and for proper cytosolic and mitochondrial Ca^2+^ dynamics upon wounding. Our work reveals a novel role for mitochondrial fission in regulating Ca^2+^ and actin dynamics during epithelial repair.

## Results & Discussion

### Mitochondrial dynamics proteins are required for wound healing

To test whether the mitochondrial dynamics machinery (Fig. 1 A) is required for epithelial repair, we performed a previously described wounding assay in the *Drosophila* embryonic epidermis (Campos et al., 2010). We laser-wounded late-stage embryos bearing wild-type and mutant alleles of mitochondrial dynamics proteins and assessed the wound healing phenotype by the percentage of non-healing wounds.

**Figure 1.**
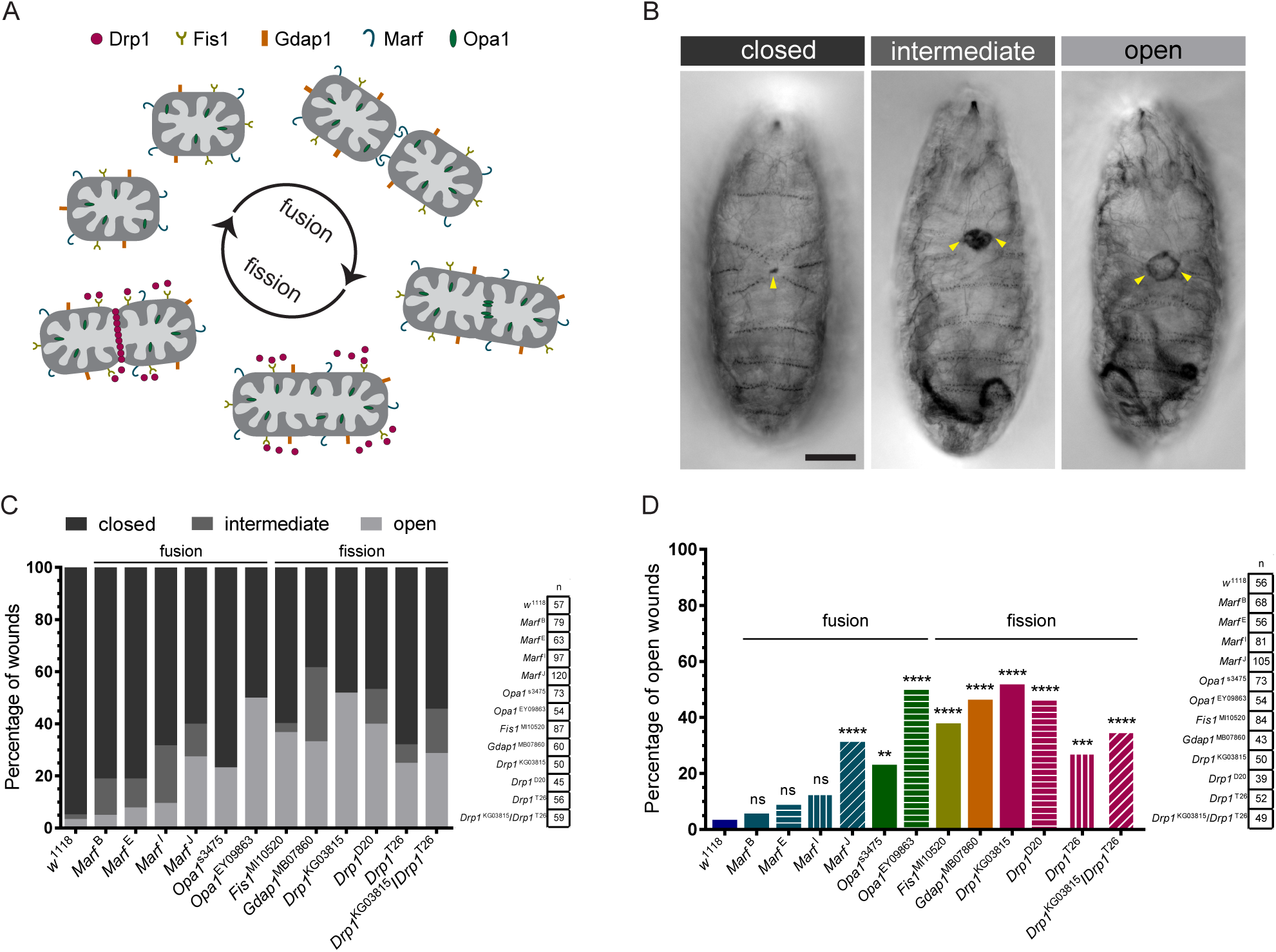
Mitochondrial dynamics proteins are required for wound healing. **(A)** Scheme of the proteins involved in mitochondrial dynamics used in the wounding assay screen. **(B)** Representative images of hatching larvae, 16 h after wounding, showing the three observed wound phenotypes: closed, intermediate and open. Closed wounds present a small scab, while open wounds show a ring of melanization around the hole. Intermediate wounds have more melanization than closed and open wounds but not a clear hole. Arrowheads point to the wound. Scale bar = 200 µm. **(C)** Graph of percentage of closed, intermediate and open wounds in controls (*w*^1118^) and mutant alleles for mitochondrial dynamics proteins. **(D)** Graph of percentage of open wounds in controls and mutant alleles for mitochondrial dynamics proteins. Regarding fusion, both *Opa1* alleles showed increased percentage of open wounds compared to controls; for *Marf*, only the *Marf*^J^ mutation shows significantly increased percentage of open wounds compared to controls. All the tested fission genes showed higher percentage of open wounds compared to controls. Fisher’s exact test was used to test for significant differences between groups. ns – not significant (P > 0.05), ** P = 0.0020, *** P = 0.0008, **** P < 0.0001. The number of embryos for each condition is shown in C and D.

Figure 1 A shows a scheme of mitochondrial dynamics with all the tested proteins represented. Regarding fusion, we tested four Mitochondrial assembly regulatory factor (*Marf*, a *Drosophila* Mitofusin homolog) alleles (*Marf* ^B^, *Marf* ^E^, *Marf* ^I^ and *Marf* ^J^) and two *Opa1* alleles (*Opa1*^s3475^ and *Opa1*^EY09863^). Concerning mitochondrial fission, we tested three *Drp1* alleles (*Drp1*^KG03815^, *Drp1*^*D20*,^ *Drp1*^T26^) and a heteroallelic combination (*Drp1*^KG03815^ / *Drp1*^T26^). We also tested other fission regulators: Fission protein 1 (Fis1), that acts as a receptor for Drp1 at the OMM (Losón et al., 2013), and Ganglioside-induced differentiation associated protein 1 (GDAP1), whose function in not well understood (Huber et al., 2013).

We observed three types of wound closure phenotypes: open, intermediate and closed wounds (Fig. 1 B). Closed wounds are identifiable by a small melanized spot. Open wounds show a melanized ring around the hole. In the intermediate phenotype, melanization occurs in a large circular area but a clear hole is absent, making it uncertain whether the wound is open or closed. Control embryos (*w*^1118^) have an outstanding capacity of epithelial repair, as 94.7% of the wounds are closed (Fig. 1 C). Mutations in either mitochondrial fission or fusion genes increased the frequency of open and intermediate wounds (Fig. 1 C).

As it is unclear whether the intermediate wounds represent a closure impairment or just a melanization defect, we excluded these wounds from the statistical analysis of the wound healing phenotype. For all mitochondrial fission genes, the percentage of open wounds was significantly higher than in controls (Fig. 1 D). Regarding mitochondrial fusion, from the four tested *Marf* alleles, only *Marf*^J^ showed increased percentage of open wounds compared to controls. *Opa1* mutants showed a significant wound closure phenotype (Fig. 1 D).

As we observed wound closure defects for mutated versions of both fusion and fission proteins, these data suggest that the regulation of mitochondrial dynamics is necessary for wound healing.

### *Drp1* mutants show delayed wound healing

Mitochondrial fission mutants showed a more consistent wound healing phenotype than fusion mutants. Therefore, we further explored the role of mitochondrial fission in epithelial repair by focusing on *Drp1* mutants. To analyze mitochondrial morphology, we used embryos expressing mitochondria [*EYFP::mito* (Lajeunesse et al., 2004)] and membrane [*PLCγPH::ChFP* (Herszterg et al., 2013)] markers. *Drp1* embryos showed longer and more interconnected mitochondria, compared to controls (Fig. S1 A-Ai, B-Bi). Mitochondrial morphology quantification confirmed that the number of branches and mitochondrial length (Fig. S1 C, D) were higher in *Drp1* compared to controls. Consistent with what has been observed in younger embryos (Macchi et al., 2013) and other *Drosophila* tissues (Sandoval et al., 2014; Verstreken et al., 2005), our results show that Drp1 regulates mitochondrial fission in the *Drosophila* embryonic epidermis.

To understand the role of Drp1 in wound healing, we used spinning-disk microscopy to image control and *Drp1* embryos expressing *GFP::Moesin* (Kiehart et al., 2000), an F-actin marker, and followed the dynamics of closure (Movie 1). Control embryos accumulate F-actin at the wound edge (Fig. 2 A) and the wound area progressively decreases until the hole is closed (Fig. 2 A-Aiii, F). Although the initial area was similar in both conditions (Fig. 2 D), *Drp1* wounds took on average 128±34 minutes (min) to close, significantly longer than controls (56±17 min) (Fig. 2 E). In milder cases, *Drp1* wounds closed at a slower rate (Fig. 2 B-Biv, F). In other cases (3 out of 13 *Drp1* embryos), the phenotype was stronger: although the wound contracted for about 40 minutes post-wounding (mpw), its area began to increase again until 120-130 mpw (Fig. 2 C-Civ, F). After this expansion phase, wounds contracted again, and in one case it was almost closed by the end of imaging (Fig. 2 Cv, F). We quantified the wound area of control and *Drp1* mutants in the first 30 mpw and found significant differences in the first minutes after wounding (4 mpw and 10 mpw for mild and strong conditions, respectively) (Fig. 2 G).

**Figure 2.**
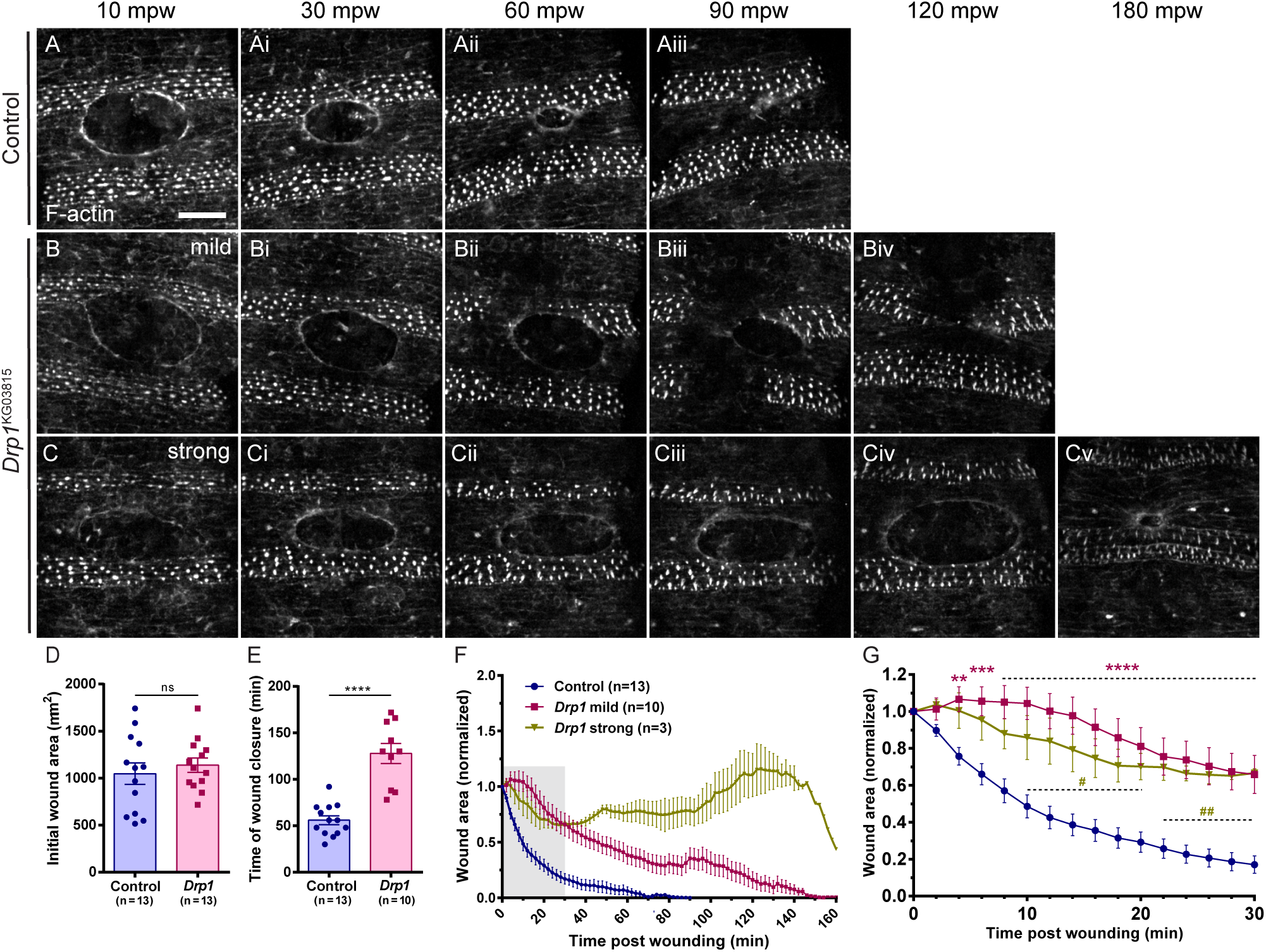
*Drp1* embryos show delayed wound healing. (A-Aiii, B-Biv, C-Cv) Maximum Z projections of the epidermis of control (A-Aiii), *Drp1* mild (B-Biv) and *Drp1* strong (C-Cv) embryos expressing an F-actin marker (*GFP::Moesin*) during wound closure. In *Drp1* mild mutants (B-Biv) wounds close slower than in controls (compare B-Biv with A-Aiii). In *Drp1* strong mutants (C-Cv), although the wound contracts in the first 30-40mpw (compare C with Ci), it then starts to expand (Cii-Civ). Later on, the wound contracts again and by 180 mpw it is almost closed (Cv). Scale bar = 20 µm. **(D)** Graph of average initial wound area in control and *Drp1* embryos (strong and mild). **(E)** Graph of wound closure time in control and *Drp1* embryos. Although the initial wound area of control and *Drp1* was similar (D), *Drp1* embryos took longer to close their wounds (E). Unpaired t test with Welch’s correction was performed to test for significant differences between groups in D and E. ns – not significant (P > 0.05), **** P ≤ 0.0001. **(F)** Graph of average wound area in control, *Drp1* mild and *Drp1* strong mutants over time. *Drp1* mild wounds close slower than controls. *Drp1* strong wounds initially contract but start to expand after 40 mpw. At 120-130 mpw wounds start to contract again. **(G)** Graph of average wound area in control, *Drp1* mild and *Drp1* strong mutants in the first 30 mpw, corresponding to the grey region in F. Significant differences between control and *Drp1* start at 4 mpw in *Drp1* mild mutants and at 10 mpw in *Drp1* strong mutants. A two-way ANOVA with a Tukey correction for multiple comparisons was used to test for significant differences between groups in G. Asterisks (*) refer to control and *Drp1* mild comparisons. Number signs (#) refer to control and *Drp1* strong comparisons. Dashed lines depict an interval of points in which the comparison between groups gives the same degree of statistical significance, given by the symbols above. # - P ≤ 0.05, ** or ## - P ≤ 0.01, *** P ≤ 0.001, **** P ≤ 0.0001. Error bars represent SEM. Number of embryos per condition is shown in each graph.

Our results show that Drp1 loss of function impairs wound closure dynamics, suggesting that mitochondrial fission is necessary for wound healing regulation.

### *Drp1* mutants have F-actin defects during wound closure

Although cells can compensate for the loss of the actomyosin cable (Ducuing and Vincent, 2016), this structure is one of the main driving forces for wound healing (Zulueta-Coarasa and Fernandez-Gonzalez, 2017). Therefore, we checked whether the wound healing phenotype in *Drp1* embryos was associated with actomyosin cable defects. We imaged control and *Drp1* embryos expressing *GFP::Moesin* (Kiehart et al., 2000) and *Zip::GFP* (Lye et al., 2014) to compare their F-actin and Myosin levels, respectively.

*Drp1* mutant embryos accumulated both F-actin (Fig. 3 B-Biii) and Myosin (Fig. 3 D-Diii) at the wound edge. However, F-actin levels were lower when compared to controls (Fig. 3 E). We found no significant differences in Myosin levels between *Drp1* and control embryos (Fig. 3 F). These results suggest that the wound healing phenotype in *Drp1* mutants is caused by defects in F-actin but not in Myosin levels.

**Figure 3.**
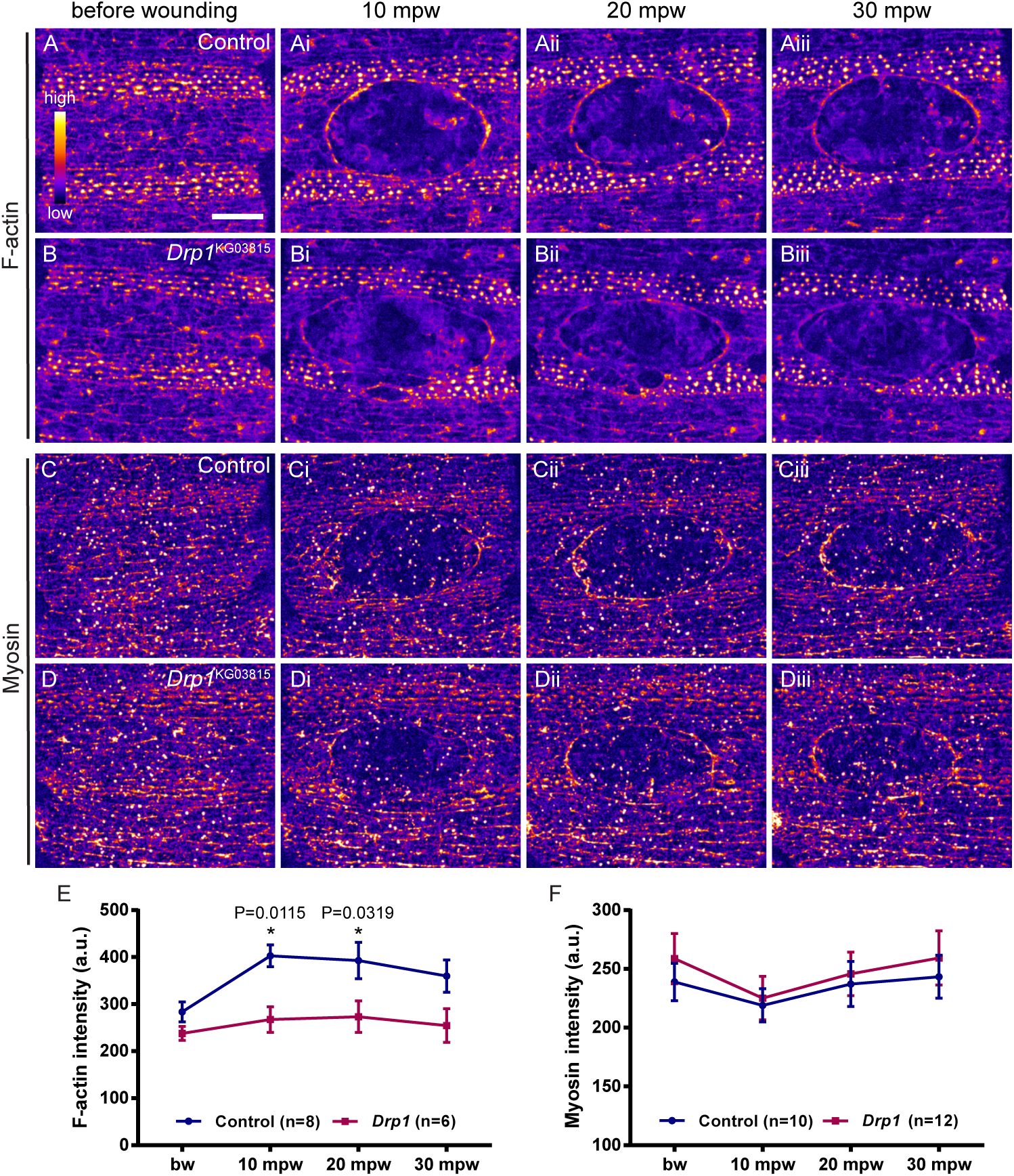
*Drp1* mutants show actin defects during wound closure. (A-Aiii, B-Biii, C-Ciii, D-iii) Maximum Z projections of the epidermis of control (A-Aiii, C-Ciii) and *Drp1* (B-Biii, D-Diii) embryos expressing an F-actin (*GFP::Moesin*) (A-Aiii, B-Biii) and a Myosin (*Zip::GFP*) (C-Ciii, D-Diii) marker before and after wounding. Images are pseudo-colored with a gradient of fluorescence intensity, ranging from blue (low) to yellow (high). Although no differences are evident before wounding (A, B), *Drp1* embryos accumulate less F-actin at the wound edge than controls (compare Bi-Biii with Ai-Aiii). Myosin accumulation at the wound edge seems similar between control and *Drp1* embryos (C-Ciii, D-Diii). Scale bar = 20 µm. **(E)** Graph of average F-actin intensity at the cell cortex before wounding and at the wound edge. F-actin levels are significantly reduced in *Drp1* mutants at 10 and 20 mpw. **(F)** Graph of average Myosin intensity at the cell cortex before wounding and at the wound edge. No significant differences were found between control and *Drp1*. A two-way ANOVA with a Sidak correction for multiple comparisons was used to test for significant differences between groups in E and F. Only significant differences (P ≤ 0.05) are represented. Error bars represent SEM. Number of embryos per condition is shown in each graph. a.u. - arbitrary units.

The formation of the actomyosin cable depends on remodeling of the Adherens Junctions (AJs) (Abreu-Blanco et al., 2012; Carvalho et al., 2014; Hunter et al., 2015; Matsubayashi et al., 2015). After wounding, the AJ protein E-cadherin (E-cad) is downregulated at the cell boundaries facing the wound, remaining only at the lateral junctions of leading-edge cells. To test whether the F-actin defects observed in *Drp1* embryos were associated with E-cad remodeling defects, we imaged control and *Drp1* mutant embryos expressing *ubi-E-cad::GFP* (Oda and Tsukita, 1999) and *mCherry::Moesin* (Millard and Martin, 2008) before and upon wounding. We observed no significant differences between E-cad levels of control and *Drp1* embryos, either before or after wounding (Fig. S2).

In summary, we propose that Drp1 regulates F-actin dynamics during wound closure, independently of AJs remodeling.

### *Drp1* mutants have altered cytosolic and mitochondrial calcium dynamics

The first signal to be detected upon wounding is an intracellular Ca^2+^ burst (Antunes et al., 2013; Razzell et al., 2013; Sammak et al., 1997; Xu and Chisholm, 2011). This Ca^2+^ increase regulates many wound healing steps, including actomyosin cable formation (Antunes et al., 2013; Xu and Chisholm, 2011). Mitochondria are known regulators of Ca^2+^ homeostasis (Finkel et al., 2015; Giorgi et al., 2008; Rizzuto et al., 2012), so we asked whether *Drp1* F-actin defects could be due to impaired Ca^2+^ dynamics.

We imaged embryos expressing the *GCaMP6f* Ca^2+^ sensor (Chen et al., 2013) and measured Ca^2+^ levels before and upon wounding (Movie 2). As previously described (Razzell et al., 2013), wounding induces a dramatic and transient increase in cytosolic Ca^2+^ (cytCa^2+^) levels in some cells around the wound (Fig. 4 Ai-Aiii. In *Drp1* embryos, the cytCa^2+^ burst was less pronounced than in controls (Fig. 4, B-Biii, C). Moreover, the area of Ca^2+^ increase was significantly reduced in *Drp1* compared to controls (Fig. 4 D), suggesting that impairing Drp1 function affects not only the Ca^2+^ levels but also the intercellular Ca^2+^ propagation.

**Figure 4.**
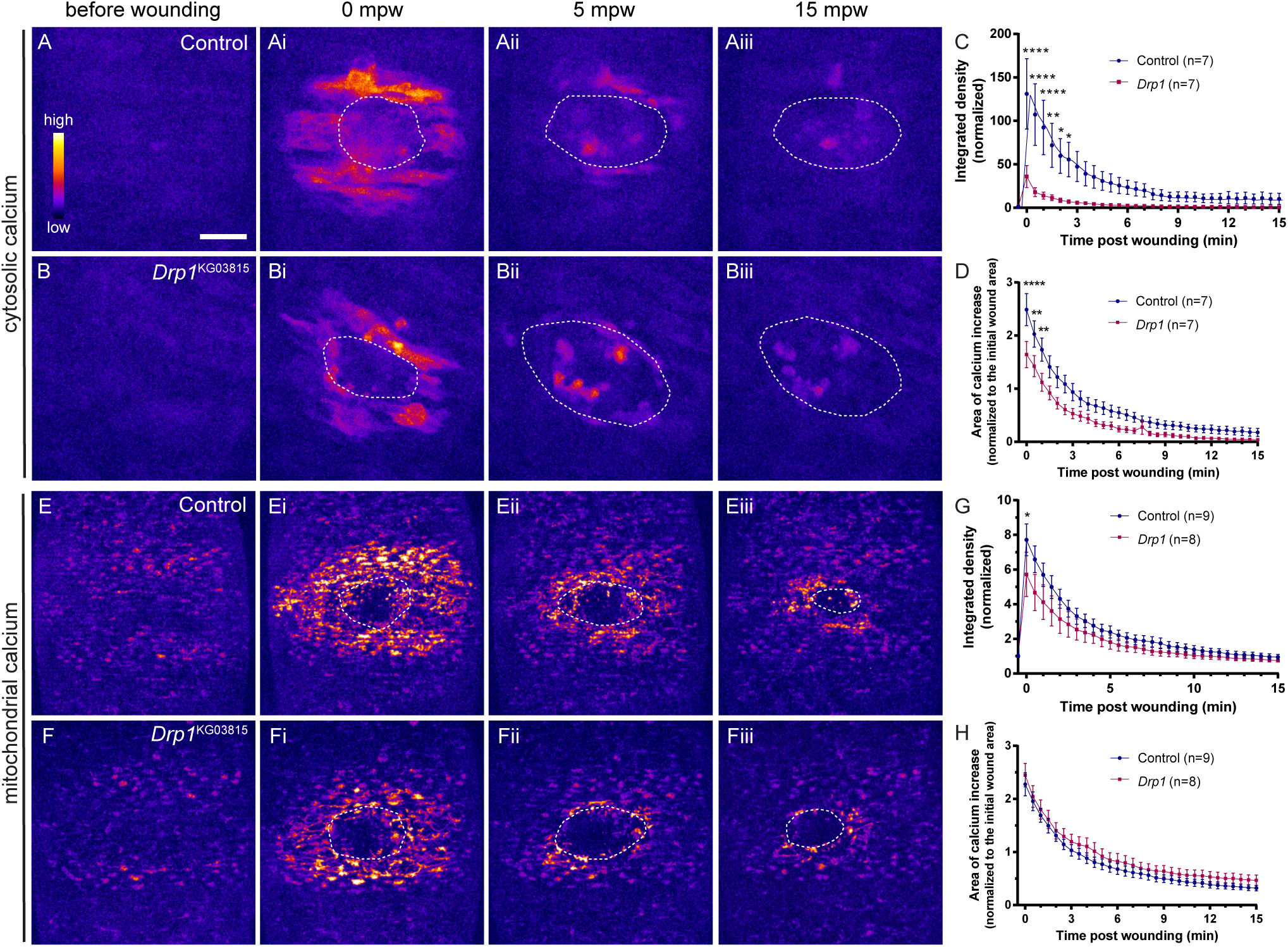
*Drp1* embryos show altered cytosolic and mitochondrial Ca^2+^ dynamics. (A-Aiii, B-Biii) Maximum Z projections of the epidermis of control (A-Aiii) and *Drp1* (B-Biii) embryos expressing a cytosolic Ca^2+^ sensor (*GCaMP6f*) before and after wounding. Both control and *Drp1* cells around the wound dramatically increase cytosolic Ca^2+^ levels immediately upon wounding (Ai, Bi). Intensity returns to pre wound levels after 15 min (Aiii, Biii). Ca^2+^ levels and area of cells that respond to the wound are lower in *Drp1* (Bi) embryos compared to controls (Ai). **(C)** Graph of cytosolic Ca^2+^ intensity in control and *Drp1* embryos shows that cytosolic Ca^2+^ is lower in *Drp1* mutants compared to controls in the first 2.5 mpw. **(D)** Graph of average area of elevated cytosolic Ca^2+^ in controls and *Drp1* embryos shows that the Ca^2+^ burst area is lower in *Drp1* embryos compared to controls from 0 to 1 mpw. **(E-Eiii, F-Fiii)** Maximum Z projections of the epidermis of control (E-Eiii) and *Drp1* (F-Fiii) embryos expressing a mitochondrial Ca^2+^sensor (mito::*GCaMP3*) before and after wounding. Wounding triggers an increase in mitochondrial Ca^2+^ levels in both control and *Drp1* cells around the wound (Ei, Fi). **(G)** Graph of mitochondrial Ca^2+^ intensity in control and *Drp1* embryos. *Drp1* embryos have a reduced Ca^2+^ burst at 0 mpw, compared to controls. **(H)** Graph of average area of elevated mitochondrial Ca^2+^ in controls and *Drp1* embryos. No significant differences were found between control and *Drp1*. Images are pseudo-colored with a gradient of fluorescence intensity, ranging from blue (low) to yellow (high). Dashed lines show the wound boundaries. Scale bar = 20 µm. A two-way ANOVA with a Sidak correction for multiple comparisons was used to test for significant differences between groups in C, D, G and F. Only significant differences are represented: * P ≤ 0.05, ** P ≤ 0.01, **** P ≤ 0.0001. Error bars represent SEM. Number of embryos per condition is shown in each graph.

Mitochondria can uptake Ca^2+^ from the cytosol, thereby modulating cytCa^2+^ (Szabadkai and Duchen, 2008). As mitochondrial morphology influences mitochondrial Ca^2+^ (mitCa^2+^) levels (Bianchi et al., 2006; Gerencser and Adam-vizi, 2005; Szabadkai et al., 2004), we examined control and *Drp1* embryos expressing a mitochondria-targeted GCaMP3 Ca^2+^ sensor [*mito::GCaMP3* (Lutas et al., 2012)] before and upon wounding (Movie 3). Concomitantly with the cytosolic response, we observed an increase in mitCa^2+^ around the wound in both control (Fig. 4 E-Eiii) and *Drp1* (Fig. 4, F-Fiii) epidermis. A mitCa^2+^ burst has been observed in *C. elegans* wound repair (Xu and Chisholm, 2014), suggesting that, similarly to the cytCa^2+^ burst, this is a conserved response to tissue injury. Quantification of mitCa^2+^ intensity showed a reduced response upon wounding in *Drp1* embryos compared to controls (Fig. 4 G, 0 mpw). This reduction is not as dramatic as seen for cytCa^2+^, maybe due to the different sensitivity of the cytCa^2+^ sensor. No differences were found in the area of increased mitCa^2+^ (Fig. 4 H), suggesting that only the cytCa^2+^ propagation is affected.

Injury triggers Ca^2+^ influx from the extracellular environment (Antunes et al., 2013; Razzell et al., 2013; Xu and Chisholm, 2011). The elevated cytCa^2+^ levels induce Ca^2+^ release from the endoplasmic reticulum (ER) mediated by the inositol-3-phosphate (IP3) receptor (IP3R), followed by propagation of Ca^2+^ and IP3 to neighboring cells through Gap Junctions (Narciso et al., 2015; Razzell et al., 2013; Restrepo and Basler, 2016). In other cellular contexts, mitochondria localize close to the ER, forming Ca^2+^ signaling microdomains. Ca^2+^ uptake by mitochondria reduces the cytCa^2+^ levels close to the open ER channels (local cytCa^2+^), preventing their Ca^2+^-dependent inactivation. By controlling ER Ca^2+^ channels activity, mitCa^2+^ uptake affects global cytCa^2+^ (Billups and Forsythe, 2002; Rizzuto et al., 2012).

Our results lead us to speculate that, upon wounding, mitochondria remove Ca^2+^ from the ER-mitochondria microdomain, preventing IP3R inactivation and favoring Ca^2+^ release from the ER. If the Ca^2+^ buffering capacity of mitochondria is compromised, as it seems to be in *Drp1* embryos, the high local cytCa^2+^ levels could inhibit IP3R opening and reduce Ca^2+^ release, resulting in lower global cytCa^2+^ levels. This would then affect the Ca^2+^ wave propagation, as less Ca^2+^ and/or IP3 would cross Gap Junctions.

Xu and Chisholm reported that suppression of mitochondrial Ca^2+^ uptake impairs the wound closure process. MitCa^2+^ triggers mitochondrial ROS production, that in turn regulates the F-actin cytoskeleton through RHO-1 (Xu and Chisholm, 2014). In this work, we show that impairment of mitochondrial fission by Drp1 loss-of-function compromises mitochondrial and cytosolic Ca^2+^ increase upon wounding. Not many studies have addressed how Drp1 modulation affects the mitochondrial Ca^2+^ buffering capacity. Drp1 deficiency in myofibers (Favaro et al., 2019) and macrophages (Wang et al., 2017) increases mitCa^2+^ levels. Here we observe the opposite effect, suggesting that the role of Drp1 on mitCa^2+^ regulation may be context dependent. Further work is needed to understand how Drp1 regulates mitochondrial Ca^2+^ uptake in the *Drosophila* epidermis.

As it has been shown that Ca^2+^ regulates actin dynamics upon wounding (Antunes et al., 2013; Xu and Chisholm, 2011), the Ca^2+^ defects observed in *Drp1* mutants could be the cause of the defective F-actin accumulation at the wound edge and consequent wound healing impairment. Nevertheless, we cannot exclude the possibility that Drp1 regulates F-actin in a Ca^2+^-independent manner. Drp1 can directly bind F-actin (DuBoff et al., 2012; Ji et al., 2015), but how Drp1 can impact F-actin dynamics is not understood. Drp1 knockdown reduces the formation of actin protrusions and invasiveness of glioma cells (Yin et al., 2016). Moreover, Drp1 can bind to RHOA and activate the RHOA/ROCK pathway (Yin et al., 2016), known to regulate cytoskeleton dynamics (Amano et al., 2010). Future studies should investigate the behavior of F-actin regulatory proteins involved in wound healing in *Drp1* mutants to understand how Drp1 can impact on the actomyosin cable.

In conclusion, our work shows a novel role for mitochondrial dynamics in epithelial repair. In particular, mitochondrial fission is essential for the wound-induced Ca^2+^ increase and F-actin polymerization at the wound edge during epithelial repair.

## Materials and methods

### Drosophila strains and genetics

Flies were maintained at 25°C on standard *Drosophila* medium. *w*^1118^ flies were used as controls for the wounding assay. The mutant alleles used were *Marf* ^B^, *Marf* ^E^, *Marf* ^I^, *Marf* ^J^, *Opa1*^s3475^, *Opa1*^EY09863^, *Fis1*^MI10520^, *Gdap1*^MB07860^, *Drp1*^KG03815^, *Drp1*^*D20*^, *Drp1*^T26^. Information on the nature of the mutant alleles can be found on Flybase (Thurmond et al., 2019). The live reporter lines used were *sqh-EYFP::mito* (Lajeunesse et al., 2004) to mark mitochondria, *ubi-PLCγPH::ChFP* (Herszterg *et al.*, 2013) to mark the cell membrane, *sqh-GFP::Moesin* (Kiehart et al., 2000) or *UAS-mCherry::Moesin* (Millard and Martin, 2008) to mark F-actin, *Zip* ^*CPTI-100036*^::*GFP* (Lye et al., 2014) to mark Myosin II, *ubi-E-cad::GFP* (Oda and Tsukita, 1999) to mark E-cad, *UAS-GCaMP6f* (Chen et al., 2013) to mark cytosolic Ca^2+^ and *UAS-mito::GCaMP3* (Lutas et al., 2012) to mark mitochondrial Ca^2+^. UAS lines were expressed under the control of the e22c-Gal4 driver.

*ubi-PLCγPH::ChFP* and *UAS-mito::GCaMP3* were a gift from Y. Bellaïche and F. Kawasaki, respectively. *Zip* ^*CPTI-100036*^::*GFP* and *ubi-E-cad::GFP* were obtained from the Kyoto *Drosophila* Genomics and Genetic Resources Stock Center, Kyoto Institute of Technology, Kyoto, Japan. All the remaining fly lines were obtained from the Bloomington *Drosophila* Stock Center, Indiana University, Bloomington, USA.

For live imaging, the *Drp1*^KG03815^ allele was recombined with live reporter lines. Mutant alleles and recombinant lines were crossed to balancer stocks that express GFP driven by a *Twist-Gal4* driver (Halfon et al., 2002). Homozygous mutant embryos were identified by the absence of GFP fluorescence. Stage 15-16 embryos were selected by the shape of the yolk (Campos-Ortega and Hartenstein, 1985).

### Wounding assay

The wounding assay was performed as previously described (Campos et al., 2010). Fly lines were in-crossed in laying pots and embryos were collected at 25°C overnight in apple juice agar plates. Embryos were dechorionated in 50% bleach and rinsed extensively with water. Selected mutant and control embryos were mounted on double-sided tape affixed to a slide, covered with halocarbon oil 700 (Sigma-Aldrich) and a 32×32mm coverslip, and sealed with polish. A 24×24mm coverslip bridge was used between the slide and the top coverslip to avoid embryo squashing.

The embryos were wounded at 25°C by using a nitrogen laser-pumped dye laser (435 nm; Micropoint Photonic Instruments) connected to a Nikon/Andor Revolution XD spinning-disk confocal microscope with an electron-multiplying charge-coupled device (EMCCD) camera (iXon 897) using iQ software (Andor Technology) and using a 60× Plan Apochromat VC Perfect Focus System (PFS) 1.4 NA oil-immersion objective.

After wounding, the top coverslip was carefully removed and the embryos were left to recover in a humid chamber at 20°C. About 16h later, the wounded embryos were scored under a stereo microscope for closed/intermediate/open wounds.

The percentage of open wounds was calculated as the ratio of nearly hatching embryos with open wounds over the total number of wounded embryos (dead animals and intermediate wound phenotypes were excluded).

Images of representative embryos depicting open, intermediate and closed wounds were acquired using a Zeiss Axio Imager Z2 widefield system equipped with an Axiocam 506 monochromatic CCD camera, a 10x EC Plan-Neofluar 0.3 NA objective and the Zen Pro 2012 software. Individual z slices with a step size of 10 µm were acquired. Stacks were processed using the Extended Depth of Field plugin based on the complex wavelet method on Fiji (Forster et al., 2004; Schindelin et al., 2012).

### Live imaging

Live imaging was performed as described previously (Carvalho et al., 2018). Dechorionated stage 15 embryos were mounted on their ventral side on glass-bottomed culture dishes (MatTek) with embryo glue (double-sided tape diluted in heptane) and covered with halocarbon oil 27 (Sigma-Aldrich). Embryos were wounded as described above for the wounding assay except that the laser power was lower in order to inflict smaller wounds that are able to close during the imaging procedure.

Time-lapse microscopy of transgenic embryos was performed at 25°C on a Nikon/Andor Revolution XD spinning-disk confocal microscope with a 512 EMCCD camera (iXon 897) with a 60× Plan Apochromat VC PFS 1.4 NA oil-immersion objective or a 60× Plan Apochromat VC PFS 1.2 NA water-immersion objective (Nikon) and using the iQ software.

Individual z slices with a step size of 0.28 µm (Fig. 1, 2 and 3), 0.36 µm (Supp Fig. 1) or 0.5 µm (Fig. 4) were acquired for a single timepoint (Fig 1) or every 30 s (Fig 4), 2 min (Fig 2) or 10 min (Fig 3, Fig S1) for 30–160 min. For F-actin, myosin and E-cad imaging, z stacks were acquired with frame averaging of 2.

### Image analysis and quantifications

All images were processed and analyzed using Fiji (ImageJ; National Institutes of Health [NIH]; (Schindelin et al., 2012), unless stated otherwise. Z-stacks were processed to obtain maximum Z-projections.

#### Mitochondrial morphology

*EYFP::mito* z-stacks were deconvolved with Huygens Remote Manager (Scientific Volume Imaging, The Netherlands, http://svi.nl), using the Classic Maximum Likelihood Estimation (CMLE) algorithm, with Signal to Noise Ratio (SNR):15 and 30 iterations. Individual cells were manually outlined and cropped from maximum Z projections of deconvolved sqh-mito-YFP merged with *PLCγPH::ChFP* z-stacks. Mitochondrial morphology from the selected cells was quantified using MiNA (Mitochondrial Network Analysis) 2.0.0 macro for Image J (Valente et al., 2017) (https://github.com/StuartLab/MiNA), selecting a Maximum Entropy Threshold Method and Ridge Detection. The branch length mean and network branches mean output parameters for each cell were plotted. The branch length mean, which was called mitochondrial length for simplicity, is the mean length of all the lines used to represent the mitochondrial structures. The network branches mean is the mean number of attached lines used to represent each structure.

#### Wound area

GFP::Moesin Maximum Z projections were used. An ellipse was drawn along the wound edge over time, and the area was obtained using the Measure tool. For each embryo, the area was normalized relative to the initial wound area. For statistical comparisons, only the first 30 min after wounding were considered, as shortly after that wounds start to close in control embryos.

#### Fluorescence intensity measurements

To measure mitochondrial and intracellular Ca^2+^ dynamics, GCaMP-6f maximum projections were used after using a median filter (0.5 pixel). The wound area, measured from *mCherry::Moesin* maximum projections from respective embryos, was deleted from GCaMP-6f maximum projections to exclude the signal coming from cellular debris and wound-recruited hemocytes. The region of Ca^2+^ increase upon wounding was selected by applying an Intensity Threshold (Otsu). The Mean Grey Value, Area and Integrated Density (the product of Area and Mean Grey Value) were obtained using the Measure Tool, before and during wound closure. We plotted the Integrated density normalized to pre wound values and the area of Ca^2+^ increase normalized to the initial wound area.

To measure F-actin and myosin intensities at the wound edge, maximum Z projections of *mCherry::Moesin*, and *Zip::GFP* stacks were used after Rolling Ball Background Subtraction (15 pixel). The wound edge and the cortical region of epithelial cells (10 cells per embryo) before wounding were outlined using a 3-pixel-wide segmented line, and the mean grey value was obtained using the Measure tool. For F-actin quantifications, cells containing actin-rich denticle precursor structures were excluded as they mask the actin present at the cable and cell cortex.

To measure E-cad intensities, maximum Z projections of *ubi-E-cad::GFP* stacks were used. Background fluorescence was subtracted from each image. The *mCherry::Moesin* channel was used to confirm the location of the wound edge. Junctions were outlined using a 4-pixel-wide segmented line and the average intensity obtained using the Measure tool. To calculate the intensity decrease (fold change) at the wound edge, the intensity value for each wound edge junction after wounding (10 and 30 mpw) was divided by the intensity value obtained for the same junction before wounding.

#### Statistics

Statistical analysis was performed using GraphPad Prism 6.01 (GraphPad Software, La Jolla California, USA). Statistical tests, P values, sample sizes, and error bars are indicated in the respective figure legends.

## Acknowledgments

We are grateful to T. Pereira and C. Crespo for invaluable support on microscopy and data analysis; E. Sucena, G. Martins and N. Pimpão for microscopy support; the Tissue Repair and Inflammation Lab and the CEDOC Fly Community for helpful discussions and technical help; the microscopy facilities at CEDOC and IGC; the fly facility at CEDOC; the Bloomington Drosophila Stock Center (NIH P40OD018537), the Kyoto Drosophila Genomics and Genetic Resources Stock Center, Y. Bellaïche and F. Kawasaki for providing *Drosophila* lines.

This work was supported by national funds through FCT - Fundação para a Ciência e a Tecnologia, I.P., in the context of a program contract to L. Carvalho (4, 5 and 6 of article 23.° of D.L. no. 57/2016 of 29 August, as amended by Law no. 57/2017 of 19 July), PD/BD/106058/2015 to S. Ponte and PTDC/BIA-BID/29709/2017; the European Research Council (2007-StG-208631) and CONGENTO LISBOA-01-0145-FEDER-022170.

No competing interests declared

## Author contributions

Susana Ponte designed, performed and analyzed all experiments (except Fig. S2) and wrote the paper. Lara Carvalho acquired funding, helped conducting and analyzing experiments in Fig. 1 and Fig. S2, and writing the paper. Isabel Campos did preliminary experiments. Maria Gagliardi did preliminary experiments, contributed to experimental design and edited the paper. Paulo J. Oliveira co-supervised the study and revised the paper. António Jacinto acquired funding, contributed to experimental design, supervised the study, and edited the paper.

